# Larval zebrafish maintain elevation with multisensory control of posture and locomotion

**DOI:** 10.1101/2024.01.23.576760

**Authors:** Samantha N. Davis, Yunlu Zhu, David Schoppik

## Abstract

Fish actively control posture in the pitch axis (nose-up/nose-down) to counter instability and regulate their elevation in the water column. To test the hypothesis that environmental cues shape strategies fish use to control posture, we leveraged a serendipitous finding: larval zebrafish (*Danio rerio*) lose swim bladder volume and sink mildly after acute loss of lateral line hair cells. Using long-term (48 h) recordings of unrestrained swimming, we discovered that sinking larvae compensated differently depending on light conditions. In the dark, they swim more frequently with an increased nose-up posture. In contrast, larvae in the light do not swim more frequently, but do climb more often. Finally, after lateral line regeneration, larvae returned to normal buoyancy and swam comparably to control siblings. We conclude that larvae can switch postural control strategies depending on the availability of visual information. Our findings complement and extend morphological and kinematic analyses of locomotion. More broadly, by quantifying the variation in strategies our work speaks to the evolutionary substrate for different balance behaviors.

## INTRODUCTION

Actinopterygian (ray-finned) fishes adjust both posture and density to regulate their depth, allowing them to occupy a remarkable range of elevations. Postural control is dynamic (Webb, 2002), and fishes will reorient relative to gravity and their environment (Meyer et al., 1976a; Meyer et al., 1977). While most fishes maintain a dorsal-up posture (Webb, 2006), notable exceptions speak to behavioral flexibility. For example, *Syndodontus nigriventris* (Meyer et al., 1976b; Willoughby, 1976) swim ventral-up, *Aeoliscus punctulatus* (Fish and Holzman, 2019) swim vertically with their heads pointed downward, and *Pleuronectiformes* swim on their side (Platt, 1973). Moreover, reorienting posture relative to gravity permits navigation in depth (Zhu et al., 2024), complementing morphological specializations that decrease density (Alexander, 1990; Pelster, 1997). Key among these specializations is the swim bladder, a gas-filled organ used by most teleosts, especially at larval stages, to countermand sinking by reducing density (Pelster, 2009; Steen, 1970). The remarkable diversity in behavior and habitat reflects selective pressures acting, at least in part, on postural control strategies.

Understanding variability in postural behavior requires perturbations. For example, swim bladder inflation can be manipulated by removing gas, attaching weights to fish, or changing external hydrostatic pressure (Fänge, 1966; Gee and Holst, 1992; Lindsey et al., 2011; Steen, 1970). *Oncorhynchus nerka* adopt nose-up postures and increase pectoral fin movements as external hydrostatic pressure increases (Harvey, 1963; Harvey and Bothern, 1972). Comparably, larval zebrafish (*Danio rerio*) change static posture and swim frequency to compensate for oil-filled swim bladders (increased density) (Ehrlich and Schoppik, 2017a) or when swimming in 1.5% glycerol (increased buoyancy) (Ehrlich and Schoppik, 2017b); neither manipulation is compatible with long-term imaging. Taken together, the work establishes that fish change posture when challenged, but the strategies they use are underexplored.

Previous work establishes the larval zebrafish as an excellent model to investigate posture, and suggests a possible route to challenge buoyancy. First, larvae swim in short translational bouts and, due to their low Reynolds number, do not glide (McHenry and Lauder, 2005). The passive period between bouts allows direct measurements of the effects of physical forces that challenge posture (e.g. gravity) (Ehrlich and Schoppik, 2017a). Recent work measuring larval posture and locomotion from the side (Zhu et al., 2023) reveals that larvae use a combination of trunk rotations and pectoral fin-generated lift (Ehrlich and Schoppik, 2019) to navigate in depth (Zhu et al., 2024). Next, larvae inflate their swim bladders early in development to reduce density (Goolish and Okutake, 1999). Further, they sense flow using a set of superficial hair cell sensors known as the lateral line; these hair cells can be killed pharmacologically and can regenerate (Davis et al., 2020). Early perturbations (pharmacological or genetic) to the lateral line interfere with normal swim bladder inflation (Lin et al., 2023; Venuto et al., 2023), suggesting a means to vary buoyancy.

Postural control is multisensory (Mergner and Hlavacka, 1995; Peterka, 2018). Visual information can shape postural behavior both by changing arousal state (Liao, 2006), and by providing feedback (Edwards and Prilutsky, 2017). Further, vision shapes flow orientation in both individual fish (Coombs et al., 2020) and tadpoles (Brown and Simmons, 2016) and groups of fish (Chaput et al., 2023; Peterson et al., 2024; Ribeiro et al., 2022; Tidswell et al., 2024). We therefore anticipate that illumination would similarly shape the way larval zebrafish stabilize posture.

We hypothesized that, if their buoyancy changed, larval zebrafish would respond by changing posture and locomotion. Further, we proposed that the nature of these changes would depend on the availability of visual information. We first observed that after loss of the lateral line, swim bladders shrank and larvae sank slightly between swim bouts; we could thus challenge fish in the light and the dark. We discovered that larvae move more frequently in the dark, but climbed more often in the light; these changes could partially recover elevation. Finally, behavior in larvae that had regenerated their lateral line was indistinguishable from age-matched control siblings. Our results reveal that, when challenged, larval zebrafish adjust posture to maintain elevation with strategies that vary with the availability of visual information.

## MATERIALS AND METHODS

### Animals

All procedures involving larval zebrafish (*Danio rerio*) were approved by the New York University Grossman School of Medicine Institutional Animal Care & Use Committee (IACUC). Zebrafish larvae were raised at 28.5°C on a standard light-dark cycle (14 hours light, 10 hours dark) at a density of 20–50 larvae in 25–40 mL of E3 medium before 6 days post-fertilization (dpf). During experiments, larvae older than 5 dpf were fed cultured rotifers (Reed Mariculture) daily for 30 minutes.

### Zebrafish lines

Behavioral experiments were performed on the Schoppik lab’s wild-type background (AB/WIK/TU/SAT/NHGRI mix), originally characterized in Zhu et al. (2023). Imaging experiments used the *Tg(myo6b:actb1-EGFP)* transgenic line to label hair cells of the lateral line and inner ear.

### Experiments

#### Behavior and Copper Treatment

Freely moving larvae were measured using the previously described Scalable Apparatus for Measuring Posture and Locomotion (SAMPL) (Zhu et al., 2023). In brief, 6–8 larvae were placed into custom vertical chambers (25 mL volume, 50 mm x 50 mm x 10 mm) filled with E3. Chambers were placed between an infrared light source (940 nm) and a camera fit with an infrared filter (Edmund Optics). Recordings were performed from the side in a light-tight enclosure with programmable white light illumination inside the box. Lighting condition is determined using the white light, as infrared light is outside of the visible spectrum of larval zebrafish (Waalkes et al., 2024). Enclosure walls, chamber holders, camera, and infrared light module are all visible with white light illumination. The field of view (400 mm^2^) was sampled from the lower middle of the chamber at a rate of 166 frames per second.

Experiments began at 6 dpf with 24 hours of recording. At 7 dpf, larvae were either treated with 10 µM copper sulfate (CuSO_4_; Acros Organics 197722500) in E3 or untreated (transferred to E3 alone) for 90 minutes. Untreated fish were transferred and handled identically to copper-treated larvae. Larvae were then washed in E3 and returned to chambers for 24 hours of posttreatment recording. Copper sulfate treatment was repeated after 24 hours to avoid hair cell regeneration (Thomas et al., 2014). Larvae were fed for 30 minutes each day; behavior was not measured during feeding. A total of 5 experimental repeats using sibling controls (i.e., 5 clutches) were completed in the dark (N=114 total larvae per condition), and 8 in the light (N=150 total larvae per condition). Developmental stages were selected for study to be consistent with the majority of work using larval zebrafish, including previous characterizations using SAMPL (Zhu et al., 2023). Further, due to their low Reynolds number, larvae at these stages do not glide (McHenry and Lauder, 2005). Consequentially it is possible to dissociate movements that follow swimming from passive destabilization, a key feature of this study.

Extensive description of SAMPL analyses and a full Python-based analysis suite are freely available at Zhu et al. (2023). In short, swim bouts were extracted from captured epochs and aligned to peak speed. Bouts were defined as lasting 250 ms before to 200 ms after peak speed. Navigational categories were classified by bout trajectory at peak speed. Climb bouts had trajectories greater than 0.5°, dive bouts were less than −0.5°, and trajectories between −0.5° and 0.5° was categorized as “flat.” The same criteria were applied to larvae in each lighting condition and treatment group. Only daytime activity (9:00–23:00) was analyzed to minimize the impact of circadian cues.

To evaluate larval ability to maintain elevation, chambers were filled with 6 mL of 2% agarose (Invitrogen 16500-500) so that the floor of the arena was in the field of view. Larvae were split into four groups: dark control (N=36), dark copper-treated (N=31), light control (N=57), and light copper (N=57) as described above. The number of larvae resting on agarose was counted every 30 minutes for 4 hours (8 observations total) across 5 experimental repeats in the dark and 6 repeats in the light.

#### Longitudinal Experiments

Effects of regenerated hair cells were examined over two periods of time: 7–9 dpf (early) and 14–16 dpf (late). Behavior recordings and copper treatments were conducted in the same manner as described above. In addition to an untreated sibling control group (N=85), one group of larvae was treated with copper sulfate in the early phase (N=94) and another group of larvae was treated during the late phase (N=64). Between behavioral recordings, larvae were raised in 450 mL of dense rotifer solution in 2 L tanks.

#### Imaging

One hour after copper treatment, larvae were washed three times for 5 minutes with E3. Larvae were next anesthetized in MS-222 (Sigma-Aldrich E10521) then mounted dorsally (lateral line) or laterally (inner ear) in 2% low-melting point agarose in E3 for imaging. Control (N=8) and copper-treated (N=15) larvae were visualized using a confocal microscope (Zeiss LSM800) equipped with a 20x water-immersion objective (1.0 NA). Lateral line and inner ear hair cells were imaged to demonstrate hair cell survival on the surface and in the otic capsule respectively. The SO2 neuromast was identified for quantification due to its large quantity of hair cells and stereotyped location. The imaging parameters (e.g. laser power, dwell time, and PMT gain) were fixed for this experiment, and did not change across fish. Further, as treatment status could be inferred unambiguously on visualization, we did not perform this experiment blind as to treatment. The swim bladder was imaged by mounting fish dorsally and laterally and measured using the three axes of volume (anterior-posterior, dorsal-ventral, width side-to-side). All images were analyzed using Fiji/ImageJ (Schindelin et al., 2012).

#### Statistical Analysis

Sample sizes were determined with respect to extensive prior characterization of wild-type variability (Zhu et al., 2023). An experimental repeat consists of all experiments run on a single clutch of fish (i.e. siblings). As most features of raw kinematic data were not normally distributed, central tendency and variance are reported as the median and interquartile range unless otherwise noted, and differences in medians between two groups were analyzed with Mann-Whitney *U* tests. Notably, since the *U* test is a rank-sum test, and there are a fixed number of experimental repeats, we can see identical test scores and p-values across different tests.

Fitting steering and righting gains requires a large number of bouts (Zhu et al., 2023); these were calculated on data aggregated across repeats, compared using unpaired t-tests. Imaging results are reported as mean and standard deviation, and analyzed using unpaired t-tests. Effect size was estimated using Cohen’s d and listed in Supplementary Tables (Lötsch and Ultsch, 2020). Finally, two-way ANOVA was used to evaluate differences across multiple conditions (light/dark, treated/untreated) and one-way ANOVA was used to assess longitudinal data, with Tukey’s HSD post-hoc comparison to adjust for multiple comparisons. Statistical significance was set at *α* = 0.05.

#### Data sharing

All raw data and code for analysis are available at the Open Science Framework DOI 10.17605/OSF.IO/BWM2E

## RESULTS

### Larvae compensate for postural instability after lateral line ablation with increased movement frequency and nose-up posture in the dark

We used a videographic assay called the Scalable Apparatus for Measuring Posture and Locomotion, or SAMPL, to measure pitch axis posture (nose-up/nose-down) and locomotion as fish navigate in depth. In the SAMPL apparatus, larvae swim freely in a vertical chamber. Behavior is sampled in a specific field of view (400 mm^2^) located in the center of the lower half of the chamber, away from the floor and walls (Figure 1A). Larval zebrafish swim in short epochs comprised of multiple short bursts of translation, or “bouts” (Figure 1B). During the inter-bout interval (Figure 1C), larvae are largely passive, allowing us to measure postural instability. To change elevation, larvae climb or dive by adjusting bout kinematics and/or posture (Figure 1D).

**Figure 1:**
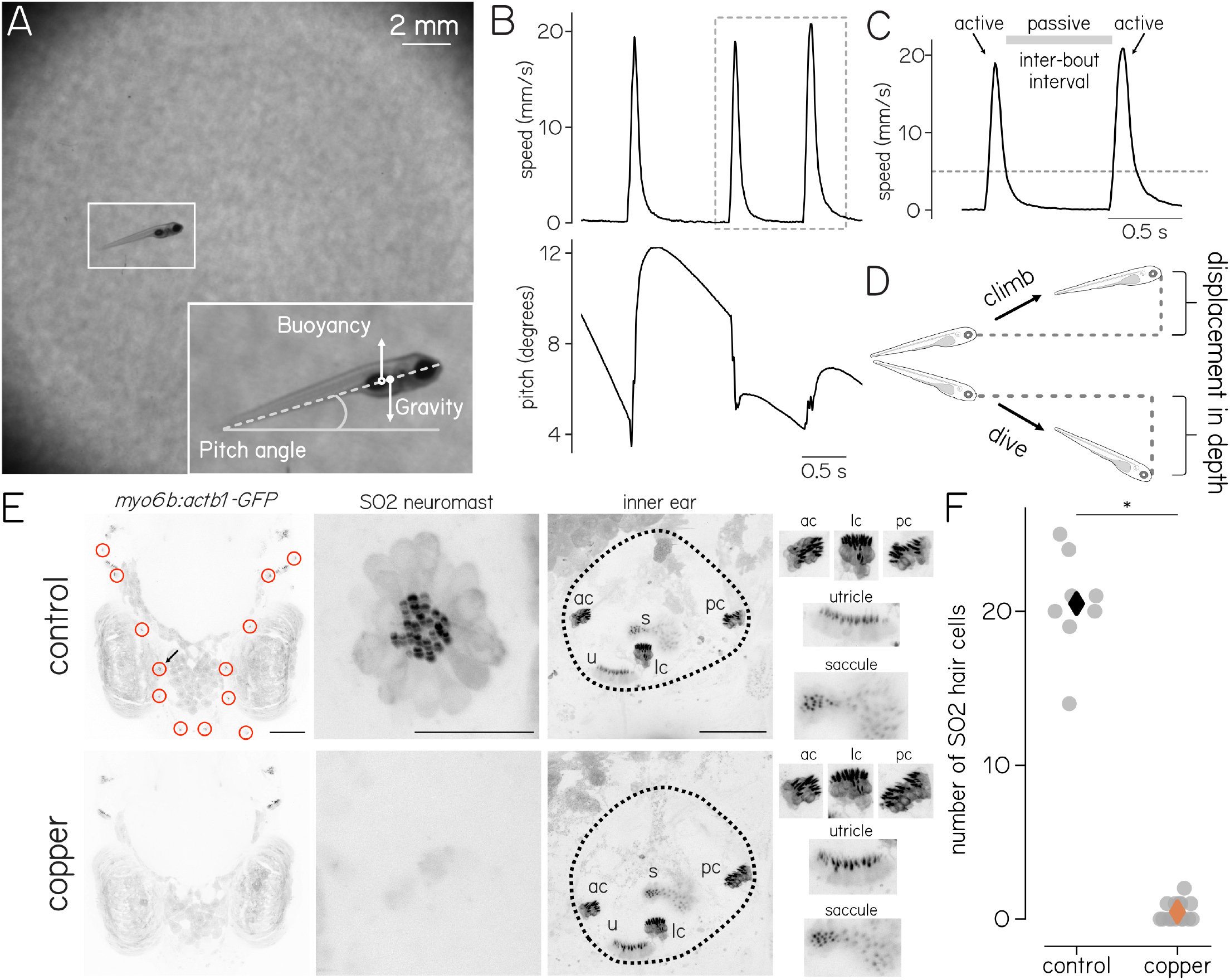
Physical forces and kinematic parameters of larval locomotion in the vertical plane. (A) Example field of view (4 cm^2^) with a 6 dpf larva. Inset shows the pitch of the larva, defined as the angle between the long axis of the body and the horizon; the gravity vector acting at the center of mass; and the buoyancy vector acting at the center of volume. (B)Representative sequence of swim bouts, plotted as speed (top) and pitch (bottom) as a function of time. Positive values for pitch are nose-up. (C)Speed trace of two swim bouts illustrating the speed threshold (dashed line; 5 mm s^-1^) that defines the active bout, as well as the passive inter-bout interval. (D)Schematic of larvae at the beginning and end of a climb and dive bout, illustrating the vertical displacement. (E)Confocal images of an untreated control fish (top) and a different copper-treated fish (bottom). Left: Dorsal view of fish head with SO2 neuromasts (red circles) and otic capsules (dotted lines) labeled. Arrow indicates a SO2 neuromast. Scale bar = 100 µm. Middle: SO2 neuromast transgenically labelled with EGFP in 7 dpf larvae. Complete loss of hair cells follows copper treatment. Scale bar = 20 µm. Right: inner ear hair cells are preserved after copper treatment. Individual end-organs are shown separately. AC, anterior crista; LC, lateral crista; PC, posterior crista; U, utricle; S, saccule. Scale bar = 100 µm. (F)Copper treatment kills lateral line hair cells. Counts and comparison made of SO2 neuromasts in 8 control and 15 copper-treated animals (p=0.0238). Grey circles and diamonds indicate individual neuromast counts and means, respectively. Unpaired t-test.

We selectively ablated lateral line hair cells by exposing larvae that were 7 days post-fertilization (dpf) to ototoxic levels (10 µM) of copper sulfate (CuSO_4_, or “copper”) for 90 minutes every 24 hours per established protocols (Hernández et al., 2006; Newton et al., 2023). Within 30 minutes, we observed that copper profoundly disrupted lateral line hair cells (Figures 1E and 1F, N = 15 copper-treated, 8 controls). In contrast, we observed no changes to inner ear vestibular hair cells that are enclosed within the otic capsule (Figure 1E). Using SAMPL, we monitored behavior of copper-treated larvae and control siblings for 24 hours before and 48 hours after copper exposure in complete darkness (Figure 2A). Before treatment, all larvae showed comparable behavior (Table S1).

**Figure 2:**
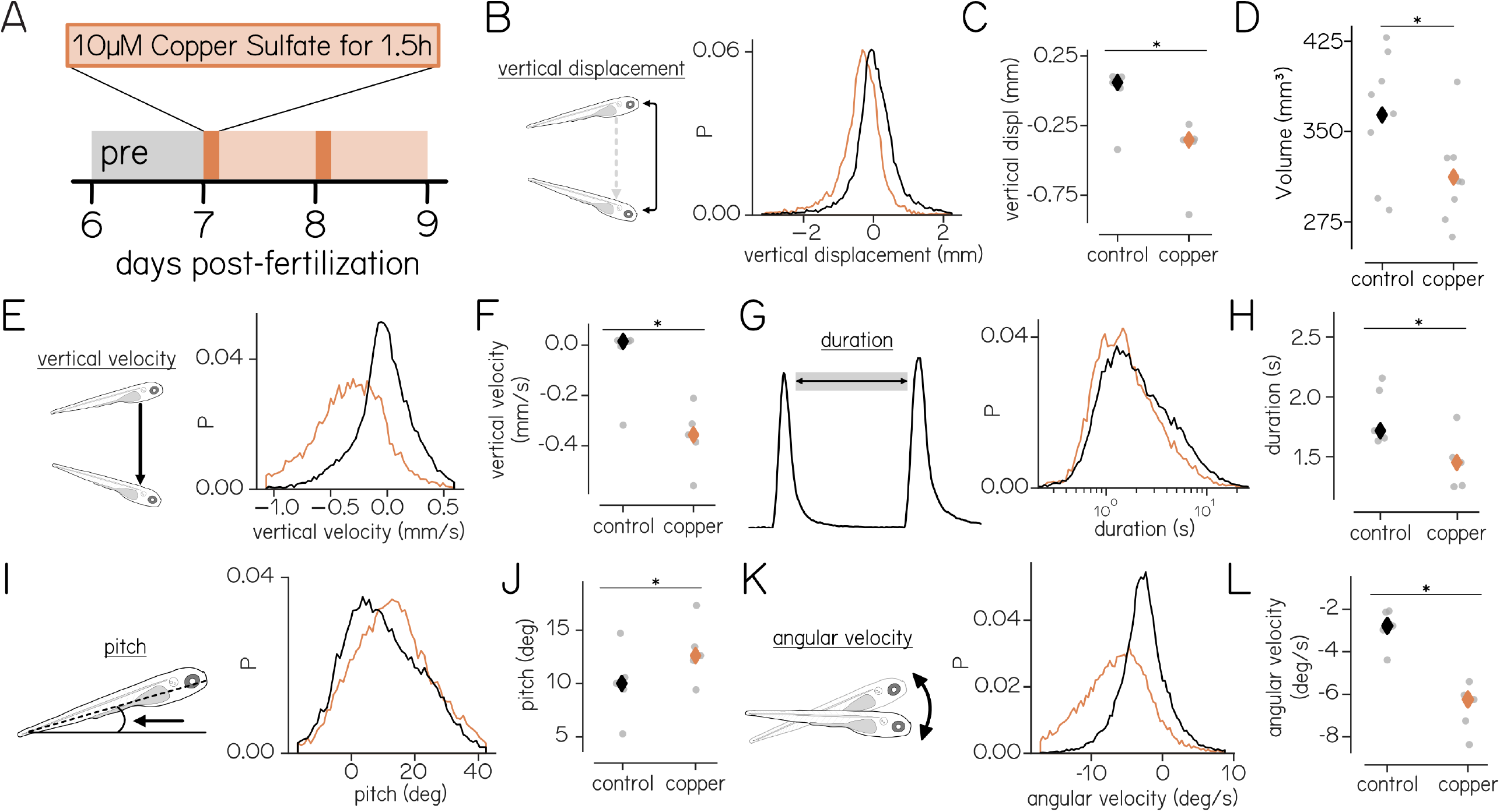
Copper treatment ablates lateral line hair cells and disrupts stability between bouts. (A)Experimental timeline indicating pre-treatment recording (grey), treatment periods (dark orange), and post-treatment recording (light orange). Full sample numbers and statistics in Table S2. (B)Schematic (left) and distribution (right) of vertical displacement between bouts. Orange are copper-treated larvae, black untreated siblings. (C)Median displacement in depth during the inter-bout interval is significantly lower in copper-treated larvae (p=0.0473). Grey circles represent experimental repeats. (D)Mean swim bladder volume is smaller in copper-treated animals (p=0.0428). See figure S1 for images and individual axis measurements. (E) Schematic and distributions of vertical velocity during the inter-bout interval. (F)Median vertical velocity between bouts is lower (i.e. downward) in copper-treated larvae (p=0.0183). (G)Schematic (left) and distribution (right) of inter-bout interval duration. (H)Median inter-bout interval duration is lower for copper-treated larvae (p=0.0300). (I)Schematic (left) and distribution (right) of inter-bout interval pitch. (J)Median inter-bout interval pitch is higher in copper-treated larvae (p=0.0473). (K)Schematic (left) and distribution (right) of angular velocity during the inter-bout interval. (L)Median inter-bout interval angular velocity is lower (i.e. nose-down rotation) in copper-treated larvae. (p=0.0060). Mann-Whitney U test. P, probability.

Copper-treated larvae sank considerably more than untreated siblings. Vertical displacement during the inter-bout interval was greater in treated larvae than untreated siblings (−0.32 [0.65] mm vs. 0.02 [0.62] mm; p=0.0473, Figures 2B and 2C, Table S2). To determine whether sinking was due to a decrease in buoyancy, swim bladder volume was calculated for each condition. Swim bladder volume was smaller in copper-treated larvae (mean [SD]: 312.22 [39.36] *µ*m^3^) compared to control animals (363.81 [52.37] *µ*m^3^; p=0.0428, figures S1 and 2D). In line with a decrease in buoyancy, copper-treated larvae sank in the vertical axis far more quickly than untreated siblings (−0.27 [0.45] mm s^-1^ vs. −0.01 [0.30] mm s^-1^; p=0.0183, Figures 2E and 2F). Sinking remained comparable across the duration of the experiment (Table S3), and can be seen during individual inter-bout intervals (Supplementary Video 1). Finally, in the inter-bout interval, treated larvae also rotated nose-down more quickly compared to untreated siblings (−6.59 [7.00]° s^-1^ vs. −2.70 [3.71]° s^-1^; p=0.0060, Figures 2K and 2L).

In response, copper-treated larvae swam more frequently and adopted a more nose-up pitch than untreated siblings. Their interbout interval duration was significantly shorter than untreated siblings (1.42 [1.15] s vs. 1.78 [2.29] s; p=0.0300, Figures 2G and 2H). In our previous work, we found that larval zebrafish with vestibular deficits do not maintain their bodies in a normal dorsal-up orientation (Zhu et al., 2024). In contrast, both treated fish and siblings maintained a largely dorsal-up posture.

However, treated fish swam with a more nose-up posture than untreated siblings (12.88 [16.80]° vs. 8.75 [17.74]°; p=0.0473, Figures 2I and 2J).

However, across experiments we observed fewer bouts in treated siblings (14699 vs. 34041, or 43%), suggesting that changes to posture and locomotion might not be completely restorative.

The sinking, increased bout frequency, increased posture, and increased angular velocity are similar to — but considerably milder than — behaviors observed in larvae that inflate their swim bladders with oil instead of air (Ehrlich and Schoppik, 2017a;Ehrlich and Schoppik, 2017b). We infer that after loss of the lateral line, larvae adjust their posture and swim frequency to partially compensate for decreased buoyancy.

### Treated larvae rely on postural changes in the dark without altering trajectory

To investigate how lateral line loss might influence bout kinematics, we examined the statistics of swim bouts after copper treatment. Swim speed was comparable between treatment conditions (copper: 15.37 [10.36] mm s^-1^ vs. control: 11.48 [6.74] mm s^-1^; p=0.0718). The average bout trajectory was also similar (copper: 13.53 [27.02]° vs. 9.77 [29.33]°; p=0.1481). We conclude that gross locomotor capacity is comparable between treated and untreated siblings.

Only climbing would countermand sinking associated with copper treatment. Using posture at the beginning of the bout, we separated bouts into climbs and dives based on their bout trajectories (Figure 3A). In the dark, the fraction of climb bouts was not significantly different (p=0.0718) between copper-treated (75 [7]%) and control siblings (66 [4]%). When climbing, we observed comparable depth changes between groups (copper: 0.53 [0.83] mm vs. control: 0.37 [0.57] mm; p=0.0718, Figure 3B). In contrast, treated larvae moved significantly further in depth during dives compared to controls, consistent with decreased buoyancy (−0.21 [0.34] mm vs. −0.16 [0.23] mm; p=0.0473, Figure 3B).

**Figure 3:**
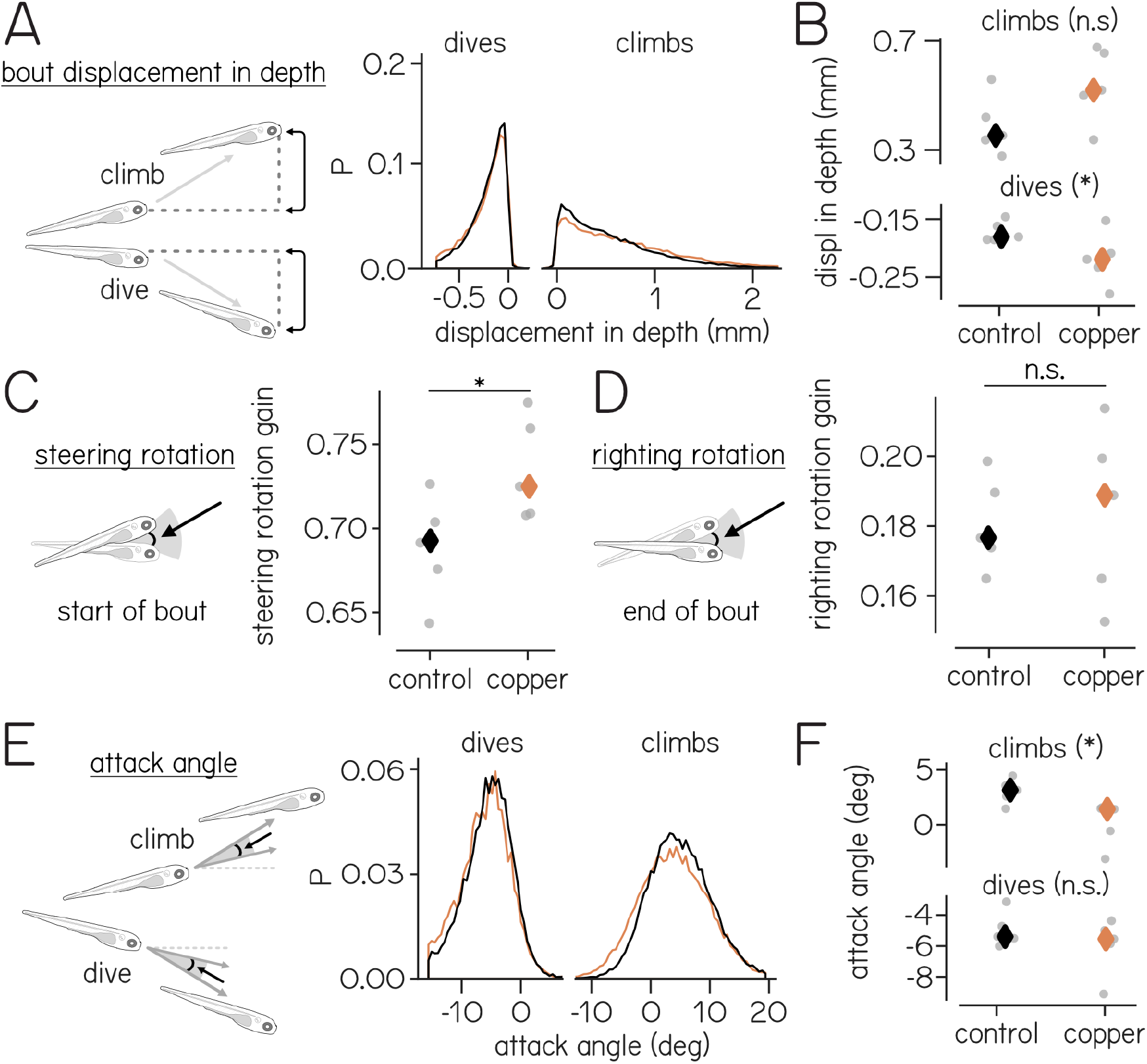
Copper-treated larvae rely on trunk rotations to produce climb bouts in the dark. (A)Schematic (left) and distribution (right) of displacement in depth during climbing and diving bouts. (B)Median displacement in depth is unchanged for climbs (top; p=0.0718) but greater for copper-treated diving bouts (bottom; p=0.0473). (C)Steering rotation gain is higher in copper-treated larvae (p=0.0367). (D)Righting rotation gain does not change between treatment groups (p=0.6281). (E)Schematic (left) and distribution (right) of attack angle during climbing and diving bouts. (F)Median attack angle is lower in copper-treated larvae for climbs (top; p=0.0300) but not dives (bottom; p=0.1050). See Table S4 for full breakdown of bouts. Mann-Whitney U test. P, probability; NS, not significant.

During climbing and diving bouts, larvae perform two distinct angular rotations using their trunk for different purposes: to change trajectory in the accelerative phase (steering), and to restore posture during the decelerative phase (righting). These rotations can be parameterized as the steering and righting gains (Ehrlich and Schoppik, 2017b; Zhu et al., 2023). Steering gain is defined as the slope of the best-fit line between posture and trajectory evaluated at the time of peak speed (Zhu et al., 2023). If the trajectory of a given bout can be explained by the posture at the time of peak speed, the steering gain would be 1. Copper-treated larvae exhibited greater steering gain than untreated siblings (0.74 [0.02] vs. 0.69 [0.02]; p=0.0367, Figure 3C). The gain of the righting rotation is defined as the slope of the best-fit line between posture at the beginning of the bout and posture change from peak speed to 100 ms after peak speed. A righting gain of 1 would indicate that the rotation perfectly restored posture. The righting gain did not differ between copper-treated and sibling larvae (0.18 [0.01] vs. 0.18 [0.02]; p=0.6281, Figure 3D).

Larval zebrafish use a second effector, their pectoral fins, to generate lift when climbing (Ehrlich and Schoppik, 2019; Zhu et al., 2023). Fin-related lift can be dissociated from steering-related changes using the difference between the predicted trajectory based on posture and the observed trajectory (“attack angle,” Figure 3E). Copper-treated larvae generated less lift, seen as lower attack angles during climbs, than sibling controls (1.14 [6.59]° vs. 3.76 [6.92]°; p=0.0300, Figure 3F). Given that larvae do not engage their fins during dives, as expected, dive bout attack angles were comparable between groups (copper: −6.27 [6.15]° vs. control: −5.25 [5.37]°; p=0.1050).

We conclude that in addition to moving more frequently and adopting a more nose-up posture, copper-treated larvae engage their trunks more when they climb. However, the average trajectory of bouts is not steeper, they do not climb more frequently, and per bout, their vertical displacement is comparable.

### Treated larvae climb more in the light

We hypothesized that visual input would influence how fish respond to lateral line hair cell loss. We therefore repeated our experiments in a standard light-dark cycle (LD) to complement data gathered in the dark (DD; Figure 4A). All measurements and comparisons across illumination conditions used data gathered during circadian day (Tables S2, S5 and S6). As expected (Prober et al., 2006), both copper-treated and untreated larvae moved more frequently in the light (Figure 4B). The inter-bout interval duration in the light was shorter for both treatment conditions (copper LD: 0.63 [0.50] s vs. DD: 1.42 [1.52] s; p=0.0010 — untreated LD: 0.69 [0.54] s vs. DD: 1.78 [2.29] s; p=0.0010). During recorded swimming epochs in the light, we observed no difference in movement frequency between copper-treated and untreated siblings (1.58 [2.00] s^-1^ vs. 1.44 [1.85] s^-1^; p=0.9000). Across experiments, we observed fewer total bouts from copper-treated fish than controls in the light (127078 vs. 182534, or 69%), though this difference was smaller than the 43% observed in the dark.

**Figure 4:**
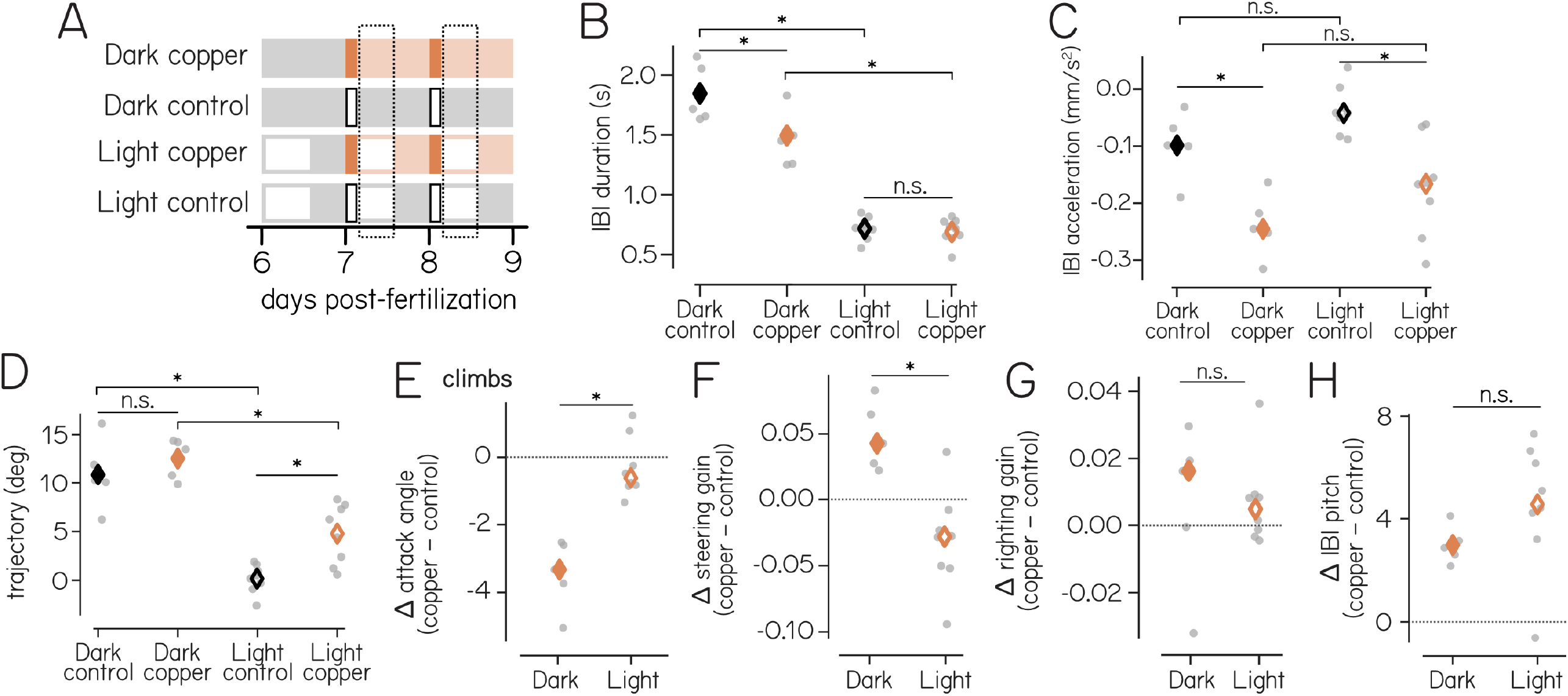
Copper-treated larvae are similarly unstable and adopt different posture and kinematic strategies in the light and in the dark. (A)Experimental timeline of each lighting condition (filled = dark, open = light) and treatment (grey = control, orange = copper). Treatment times indicated in dark orange, and dotted rectangles show time window during which behavior was compared. Full sample numbers and statistics in Tables S5 and S6. (B)Inter-bout interval (IBI) durations are longer in the dark than in the light (control: p=0.0010, copper: p=0.0010). Copper-treatment shortens IBI duration in the dark (p=0.0033) but not the light (p=0.9000). (C)Vertical acceleration during the IBI is lower (more downward) in copper-treated larvae in both the dark (p=0.0099) and the light (p=0.0017). Acceleration did not vary as a function of light for control (p=0.3608) or copper-treated larvae (p=0.2900). (D)Median swim trajectory of copper-treated larvae is higher than untreated siblings in the light (p=0.0135) but not the dark (p=0.3277). Trajectories are lower (more horizontal) in the light (control: p=0.0010, copper: p=0.0010). (E–H) Data are plotted as the difference between control and copper-treated larvae for each experimental repeat for both lighting conditions. Dotted lines at 0 indicate no difference between copper-treated and untreated siblings. P values are comparisons of the differences in the dark vs. light. (E)Differences in attack angle for climb bouts are small in the light, unlike the dark (p=0.0001). (F)Differences in steering rotation gain are higher in the dark than in the light (p=0.0017). (G)Differences in righting rotation gain are unchanged (p=0.9807). (H)Differences in IBI pitch are higher but not significantly different in the light (p=0.5123). See Table S4 for full breakdown of bouts. B–D: Two-way ANOVA with Tukey’s HSD. E–I: Mann-Whitney U test (multiple comparison p-values listed in Table S5). IBI, inter-bout interval; NS, not significant.

We next compared linear acceleration during the inter-bout interval to determine if the physical challenges that followed copper treatment were comparable in the light and the dark. Instability after lateral line ablation was comparable in both the light and dark during the inter-bout interval with a main effect of both light (p=0.0216) and treatment (p=1.5e-5) but no interaction (p=0.9176). Copper-treated larvae showed greater acceleration in the downward direction than untreated siblings in both the light and the dark (LD copper: −0.16 [0.08] mm s^-2^ vs. LD untreated −0.04 [0.03] mm s^-2^; p=0.0017 — DD copper: −0.24 [0.03] mm s^-2^ vs. DD untreated: −0.09 [0.03] mm s^-2^; p=0.0099, Figure 4C). Crucially, downward acceleration did not change between light and dark for either treatment (copper LD vs. DD: p=0.2900 — untreated LD vs. DD: p=0.3608). As linear acceleration was comparable in both light and dark regardless of treatment, we conclude that the changes after copper treatment reflect true changes to density.

Given the comparable physical challenges to maintaining elevation (i.e. increased downward acceleration), we next asked if compensatory strategies were similar after copper-treatment in the light and dark (Tables S5 and S6). Unlike in the dark, copper-treated larvae in the light climbed with steeper trajectories than untreated siblings (LD copper: 4.28 [21.82]° vs. LD untreated: −1.29 [18.31]°; p=0.0135, Figure 4D). Copper-treated larvae climbed more frequently (60 [13]%) than control larvae (46 [5]%; p=0.0013). In the light, copper-treated larvae relied less on trunk steering to climb, instead engaging their fins to produce lift. Fin engagement (attack angles) during climbs was comparable between copper-treated and untreated siblings in the light (LD control: 1.99 [4.33]° vs. LD copper: 1.71 [4.72]°; p=0.9000, Figure 4E). In the light, steering gain (trunk use) was unchanged between copper-treated larvae than untreated siblings (control: 0.75 [0.03] vs. copper: 0.72 [0.02]; p=0.2755, Figure 4F), as was righting gain (control: 0.09 [0.01] vs. copper: 0.10 [0.01]; p=0.7969, Figure 4G). However, similar to copper-treated larvae in the dark, those in the light adopted a more nose-up posture than untreated siblings (4.96 [16.71]° vs. −0.37 [14.84]°; p=0.0037, Figure 4H).

Just as in the dark, copper-treated larvae in the light changed their behavior by increasing posture. However, unlike fish in the dark, copper-treated larvae did not move more, nor did they change bout kinematics (attack angle and steering gain). Instead, they climbed more frequently, and with steeper trajectories than untreated siblings. We conclude that light can change compensatory strategies after lateral line hair cell loss.

### Light improves elevation efficiency

In both the dark and the light, we observed fewer total bouts from copper-treated larvae compared to untreated siblings (Table S4). We hypothesized that the decrease in observed bouts was due to ineffective elevation control. If so, copper-treated larvae would sit at the floor of the chamber outside of the normal field of view of the camera. We adjusted the arena to observe the chamber floor and observed larvae position in dark and light conditions (Figure 5A). There were main effects of both light condition (Figure 5B; p=3.8e-10) and treatment (p=1.4e-27), as well as an interaction between the two (p=5.8e-4). Specifically, copper-treated larvae were positioned at the bottom of the arena more often than controls in the dark (12.5 [12.3]% vs. 54.1 [50.6]% ; p=0.001) and in the light (4.8 [12.5]% vs. 33.3 [13.8]% ; p=0.001). Fewer copper-treated larvae were observed on the floor in the light relative to the dark. Importantly, light improved elevation in copper-treated larvae (p=0.001), but not sibling controls (p=0.1188).

**Figure 5:**
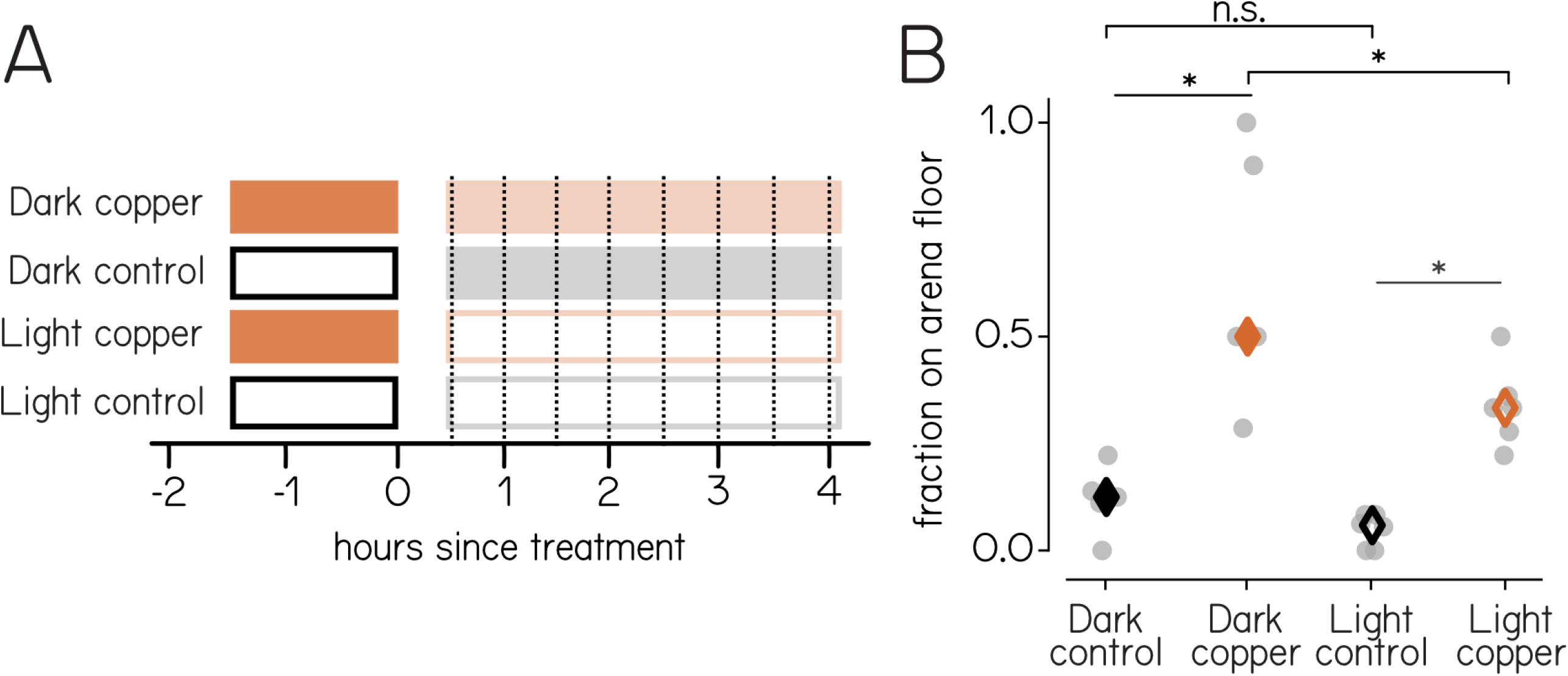
Copper-treated larvae are inefficient at maintaining elevation, but are better in the light. (A)Experimental timeline of each lighting condition (filled = dark, open = light) and treatment (grey = control, orange = copper). Treatment times indicated in dark orange, and dotted lines demonstrate observation times. Dark control (N=36), Dark copper (N=31), Light control (N=57), Light copper (N=57). (B)Fraction of larvae resting on the chamber floor is greater after lateral line ablation in both the dark (control v. copper: p=0.001) and light (control v. copper: p=0.001). Larvae in the light were less likely to rest on the floor between copper-treated groups (p=0.001) but not controls (p=0.1188). Two-way ANOVA with Tukey’s HSD.

Given the downward acceleration, fewer bouts observed, and number of larvae observed at the arena floor, we conclude that lateral line ablation compromises the ability of larvae to maintain elevation. The physical challenges faced by copper-treated larvae were comparable between the light and the dark. We therefore propose that the strategies adopted in the light are more effective at compensating for decreased buoyancy.

### Larvae recover postural stability after lateral line regeneration

We hypothesized that buoyancy would be restored following regeneration of lateral line hair cells. Larvae were treated with copper either in an early phase (7–9 dpf) or late phase (14–16 dpf) and compared to untreated control siblings (Figure 6A). Copper treatment killed lateral line hair cells similarly at both time points (Figure 2B and Figure 6B), but early-treated fish had regenerated their lateral line hair cells by 14 dpf (Figure 6B). Similar to exposure at 7 dpf, late-treated larvae sank faster than controls (−0.23 [0.46] mm s^-1^; p=0.0013) and early-treated animals (p=0.0012). In the early phase, larvae presented similarly to data presented in Figures 2 and 3. Here, we describe data collected in the late phase only.

**Figure 6:**
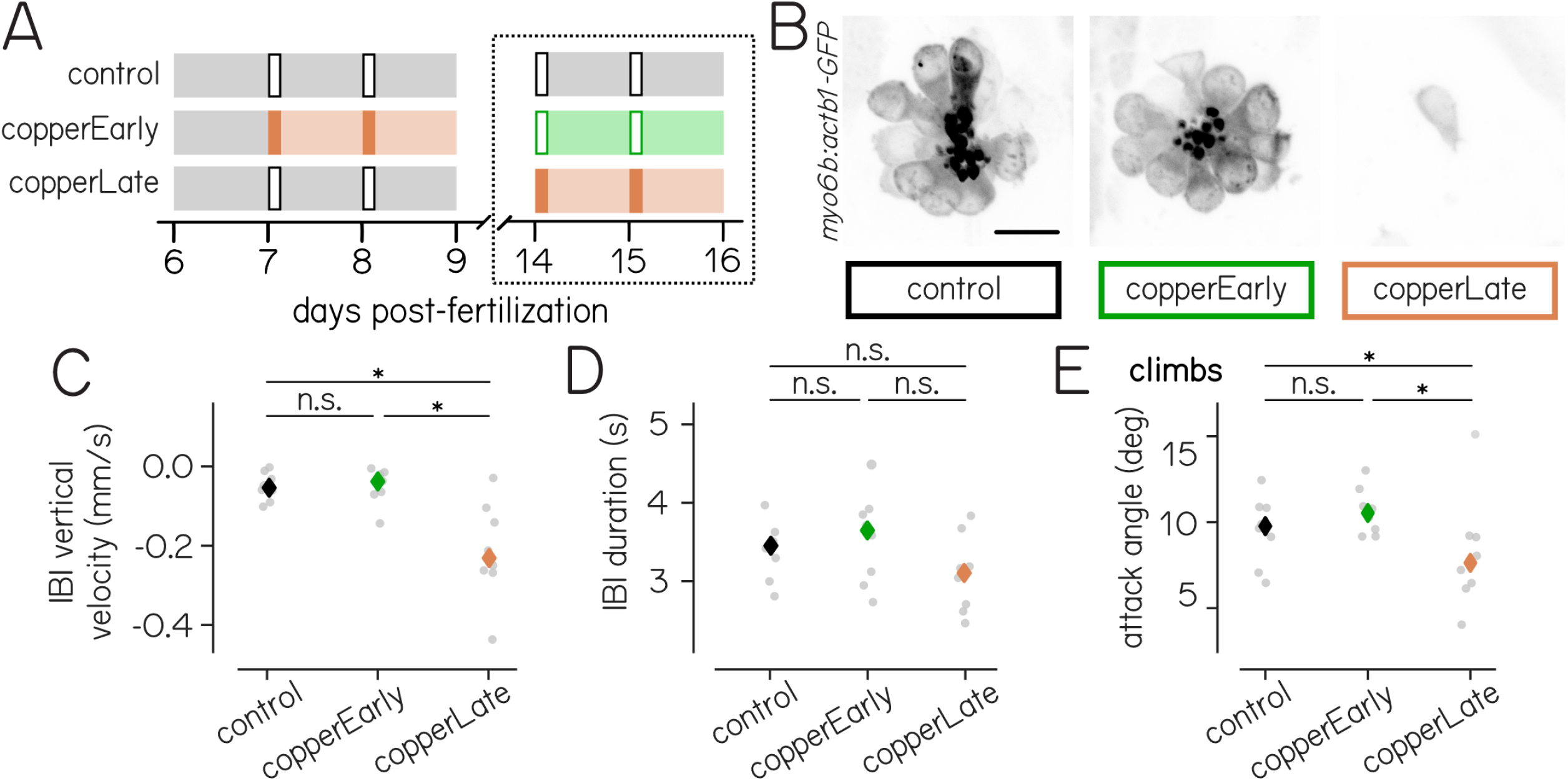
Larvae return to normal kinematics after lateral line regeneration. (A)Experimental timeline of each condition in the early (7–9 dpf) and late phase (14–16 dpf). Color indicates treatment (grey = control, orange = acute copper ablation, green = regenerated hair cells). Treatment times indicated in dark orange, dotted rectangle shows time window during which behavior was compared. Full sample numbers and statistics in Table S7. (B) Confocal imaging of SO2 neuromast hair cells at 14 dpf in representative control (left), early-treated (center), and late-treated larvae (right). Lateral line hair cells were regenerated to control levels in early-treated larvae, and similarly ablated by copper in late-treated siblings. Scale bar = 10 µm. (C)Vertical velocity during the IBI is similar between early-treated and control animals (p=0.9000), while late-treated larvae sink significantly (late v. control: p=0.0013; late v. early: p=0.0012). (D)IBI duration is similar between all groups (p=0.1832). (E)Attack angle during climb bouts is comparable between all groups (p=0.0671). (F)Peak speed is similar between early-treated and control animals (p=0.3964), but increased in late-treated larvae (late v. control: p=0.0330; late v. early: p=0.0119). See Table S4 for full breakdown of bouts. One-way ANOVA with Tukey’s HSD post-hoc comparison. disp, displacement; IBI, interbout interval; NS, not significant.

Larval buoyancy returns to control levels after hair cells regenerate. Vertical velocity between bouts was similar for early-treated larvae and controls (−0.03 [0.24] mm s^-1^ vs. −0.04 [0.31] mm s^-1^; p=0.9000, Figure 6C). Other properties of the inter-bout interval also recovered after lateral line regeneration. The inter-bout interval duration of early-treated larvae returned to control values (2.92 [3.49] s vs. 2.50 [3.69] s; p=0.7858, Figure 6D). Early-treated animals also held similar postures to untreated siblings (7.08 [18.09]° vs. 11.25 [21.51]°; p=0.5569). We conclude that that 1) lateral line regeneration resolves the positional challenges imposed by copper treatment, and 2) lateral line loss at 14 dpf similarly affects buoyancy.

Because early-treated larvae sank less and improved stability after hair cell regeneration, we asked whether their compensatory climbing behaviors persisted. Early-treated and control larvae exhibited similar attack angles (10.68 [6.21]° vs. 9.77 [6.80]°; p=0.1860; Figure 6E), which were comparable to late-treated animals (7.55 [6.11]°; p=0.0639). Larvae across all three groups performed similar fractions of climbs and dives (control: 71[7]%; early: 68 [2]%; late: 71 [20]%; p=0.6314), consistent with previous observations in the dark. However, unlike compensatory actions at 7 dpf, late-treated larvae swam faster than sibling controls (13.07 [9.91] mm s^-1^ v. 10.46 [6.56] mm s^-1^; p=0.0330; Figure 6F) and early-treated larvae (9.88 [5.89] mm s^-1^; p=0.0119), while control and early-treated larvae displayed similar speeds (p=0.3964).

Early-treated larvae swam comparably to controls after hair cell regeneration. While late-treated larvae sank like early controls, the major change to their behavior was an increase in swim speed. We conclude that hair cell regeneration is sufficient to recover buoyancy, and that stabilizing strategies can change with age.

## DISCUSSION

We hypothesized that postural control would shift to incorporate multisensory information, and investigated the ways larval zebrafish maintain elevation when buoyancy is challenged. Lateral line loss reduces the size of the swim bladder; fish become denser and sink, which triggers changes to behavior that vary in the light and the dark. Changes to behavior were only partially effective, but differentially so: treated fish were more likely to rest on the bottom of the arena, but less so in the light. Finally, sinking and compensatory changes recovered after peripheral regeneration, demonstrating transient effects of ablation. We conclude that fish have multiple strategies for maintaining elevation, and selectively engage them depending on available sensory information. The behavioral flexibility we observe could act as a substrate for selective pressures that gave rise to the vast array of postures and preferred elevations among fish.

We observed that larvae sank after copper treatment. The small increase in downward acceleration we observed was stable across clutches of fish and illumination. The most likely explanation for the changes we observe is that acute loss of the lateral line causes larvae to slightly deflate their swim bladders (Dumbarton et al., 2010), as swim bladder size is linked to density (Stewart and McHenry, 2010). Indeed, we found that swim bladder volume was lower after copper treatment. Early lateral line manipulations can induce hyperbuoyancy (Lin et al., 2023; Venuto et al., 2023); our results expand this link between the lateral line and swim bladder volume. Importantly, near-neutral buoyancy is re-established after lateral line regeneration; we propose that this follows restoration of normal swim bladder volume. We therefore infer that the decrease in swim bladder volume is reversible, supporting a proposed role for lateral line-derived signals that regulate density (Venuto et al., 2023).

The behavioral changes observed following copper exposure might suggest a disruption to the vestibular or nervous systems. Unlike larvae with constitutive vestibular impairments (Zhu et al., 2023), copper-treated fish fish maintain a dorsal-up orientation in the dark. Inner ear (vestibular) hair cells are protected by the otic capsule. While copper treatment can disrupt these cells, to do so it must be injected directly into the otic capsule (Hamling et al., 2023). Consistently, we do not observe gross changes to hair cells in the inner ear after treatment. Long-term copper exposure can induce neurotoxicity and alter swim speeds; however, our treatment has a shorter duration, lower concentration, and no effect on speed. Other compounds known to kill lateral line hair cells were not used due to effects on musculature (Han et al., 2020) and longer treatment times (Coffin et al., 2013). Taken together with the known effects of lateral line manipulations on swim bladder volume (Lin et al., 2023; Venuto et al., 2023), the preponderance of evidence supports the conclusion that the effects we see follow disruption of lateral line but not vestibular hair cells or neurotoxic effects.

Illumination-dependent findings reflect limits of behavior fish can produce when challenged with lateral line loss. First, challenged larvae swim more frequently than control siblings in the dark, with less fin engagement (lower attack angles). Neither swim frequency nor fin engagement changed in the light, presumably because in the light, both are at their ceiling. In the light larvae climbed more frequently with greater average trajectories. A similar ceiling for climb fraction and trajectory likely exists in the dark. While we restricted our analysis to comparable points in time, it would be interesting to explore these ceilings in the context of circadian day and night, particularly given diel-related elevation changes in fish (Helfman, 1986).

The strategies we observed were more effective in the light than in the dark. Evidence comes from two sources: first, we observed greater number of observed bouts (which can only happen if fish are off the bottom of the arena) in the light; second, a greater fraction of fish were observed on the bottom of the arena in the dark. The change to buoyancy, measured by downward acceleration after treatment, was comparable in the light and the dark. While there was a significant change in vertical displacement per climb bout in the light, the effect size was quite small. Instead, we propose that the improvement in the light reflects an increased fraction of climbs that, together, allows for elevation changes. Larval zebrafish sequence climbs and dives to change elevation when navigating in depth (Zhu et al., 2024). Given that illumination can drive changes in elevation (Hedenström et al., 2022; Nilsson et al., 2018), we propose that a similar mechanism explains improved control of elevation we observed in the light. One possibility is that light signals are used for locomotive control by cuing verticality, similar to insects (Fabian et al., 2024), but the mechanisms of light-mediated locomotor control remain unknown.

We propose that the changes we observe when larvae are challenged might reflect differential engagement of neural circuits responsible for postural set point, swim frequency, and fin control. Two likely, but not mutually exclusive, substrates for these behaviors are the cerebellum and the interstitial nucleus of Cajal, also called the nucleus of the medial longitudinal fasciculus (INC/nMLF). First, the cerebellum is a hub of sensory integration (Ishikawa et al., 2015), modulating vestibular function (Auer et al., 2023; Manzoni, 2007) and motor activity (Cullen, 2023; Manzoni, 2007). The cerebellum has been reported to modulate fin-body coordination (Auer et al., 2023; Ehrlich and Schoppik, 2019), a key difference in the strategies observed. Next, the INC/nMLF is likely engaged during different compensatory tactics given its activity during various swim modes (Sankrithi and O’Malley, 2010) and involvement in posture and navigation (Sugioka et al., 2023; Zhu et al., 2024). The INC/nMLF is responsive to light, head taps (Sankrithi and O’Malley, 2010), body tilts (Sugioka et al., 2023), and horizontal visual motion (Naumann et al., 2016). Future work differentiating patterns of neuronal activity following body tilts (Hamling et al., 2022) in the light and dark will speak to the substrates responsible for behavioral strategies.

How might light affect control of elevation? Visual cues during sinking might induce reflexive behavior to maintain elevation. Such stabilizing optomotor responses are well-characterized in the horizontal (yaw) plane (Clark, 1981), with well-defined neural substrates that transform optic flow into swim kinematics (Dehmelt et al., 2021; Matsuda and Kubo, 2021; Orger et al., 2008). While optomotor stabilization in other planes (pitch, roll) have not been reported, rotations in both axes can induce reflexive optokinetic responses that stabilize gaze (Bianco et al., 2012; Burkhardt et al., 2025). If vertical visual stimuli can similarly elicit compensatory swimming, the increased sinking following perturbation of the lateral line would act to enhance the strength of the optic flow cue. Complementarily, light provides both a directional cue that can modulate posture through the dorsal light reflex(von Holst, 1935), retina-independent vertical phototaxis (Fernandes et al., 2012), and generalized arousal (Liao, 2006). By establishing that strategies for postural control change as a function of illumination, our work opens the door to investigate the contribution of different reflexes on elevation control.

Dealing with gravity is a biomechanical challenge for all animals. Without a substrate, maintaining elevation becomes vital for aquatic and airborne creatures. We discovered that larval zebrafish become denser and sink after acute lateral line loss, enabling comparisons of compensatory strategies with and without light. Larvae correct for destabilization by climbing using a combination of postural and kinematic adjustments. As hypothesized, these mechanisms diverged when visual information was available into distinct strategies, with different efficacy. Our work reveals separate behaviors to accomplish the same purpose — maintaining elevation — framing future investigation into how different sensory inputs and locomotor actions regulate position.

Different behavioral strategies can serve as a substrate for natural selection (Brown and Wilson, 1956). Empirically, orientation varies across Actinopterygian species (Fish and Holzman, 2019; Meyer et al., 1976b; Platt, 1973; Webb, 2006; Willoughby, 1976), speaking to past pressures that acted on the behavioral strategies that regulate posture. Understanding how animals change behavior when challenged therefore extends existing studies into the morphological (Drucker, 2002) and locomotor specializations (Fish and Lauder, 2006; Tytell et al., 2010) that are well-suited to different environmental niches.

## ACKNOWLEDGMENTS

Research was supported by the National Institute on Deafness and Communication Disorders of the National Institutes of Health under award numbers R01DC017489, the National Institute of Neurological Disorders and Stroke under award number R61NS125280, by the Leon Levy Foundation (YZ), the Rainwater Charitable Foundation (YZ) and by the Irma T. Hirschl/Monique Weill-Caulier Trust (DS). The authors would like to thank Hannah Gelnaw for assistance with fish care, Lavinia Sheets and lab for sharing the *Tg(myo6b:actb1-EGFP)* line, Dena Goldblatt and other members of the Schoppik and Nagel labs for their valuable feedback and discussions.

## AUTHOR CONTRIBUTIONS

Conceptualization: YZ; Methodology: YZ, SD; Investigation: SD, YZ; Visualization: SD; Writing: DS, SD; Editing: SD, DS; Funding Acquisition: DS, YZ; Supervision: DS.

## SUPPLEMENTAL VIDEO

**Video S1**

Video S1. Real-time recording of control and copper-treated larvae after treatment. Notably, copper-treated larvae sink between bouts, while controls maintain their position in depth. Related to Figure 2.

**Figure S1:**
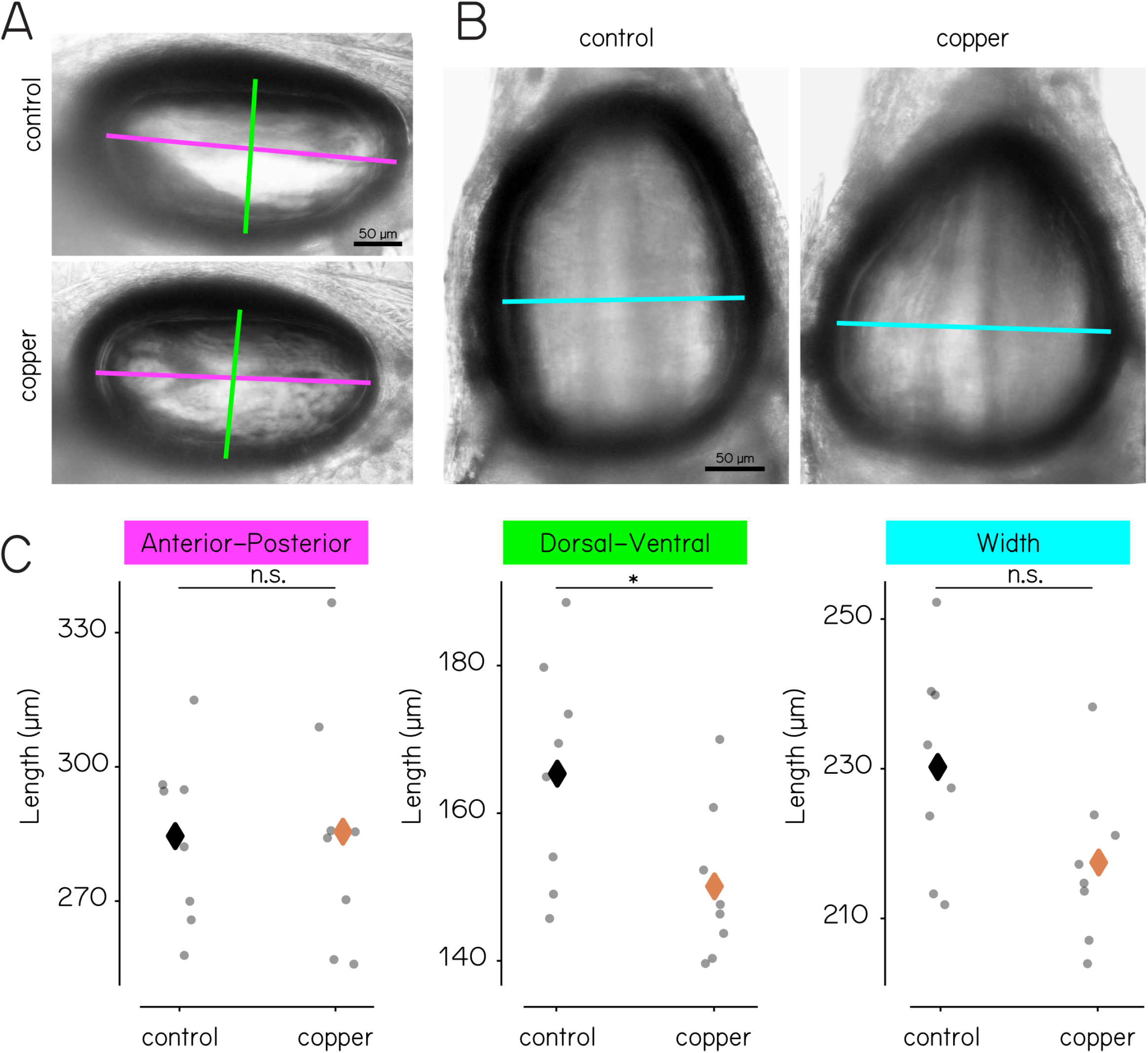
Swim bladder size decreases in the dorsoventral axis after copper treatment. (A)Example lateral-mount images of swim bladders from a control larva (top) and a different copper-treated larva (bottom) at 7 dpf. Pink axis = anterior-posterior, green axis = dorsoventral. (B)Example dorsal-mount images of swim bladders from control (left) and copper-treated (right) larvae at 7 dpf. Blue axis = width. (C)Left: A-P axis length does not change with copper treatment (p=0.9341). Middle: D-V axis length decreases with copper treatment (p=0.0355). Right: Width does not change with copper treatment (p=0.0593). Unpaired t-test.

**Table S1:**
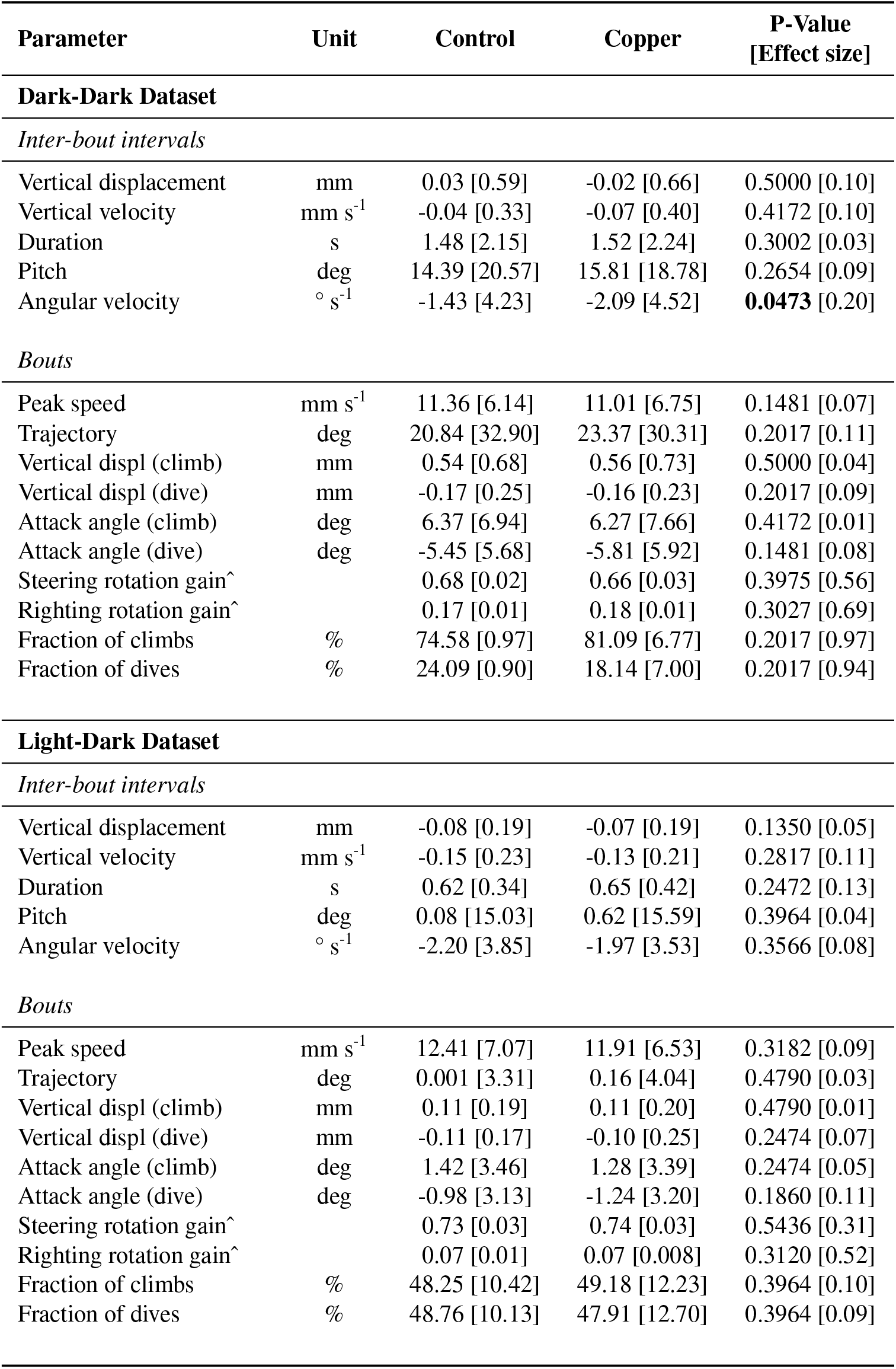
Pre-treatment results for dark-dark and light-dark datasets. Median [interquartile range (IQR)] are listed for non-normal distributions of raw data. P-value calculated using the Mann-Whitney U test (*α* = 0.05), and effect size calculated using Cohen’s d. Bold values indicate significance. ^Steering and righting rotation gain p-value calculated using unpaired t-test (mean and standard deviation listed).

**Table S2:** Effects of lateral line ablation in the dark. Median [IQR] used for non-normal distributions of raw data. N = 114 control larvae, 114 copper-treated larvae. P-value calculated using Mann-Whitney U test (*α* = 0.05), and effect size calculated using Cohen’s d. Bold values indicate significance. ^Steering and righting rotation gain p-value calculated using unpaired t-test (mean and standard deviation listed).

| Parameter | Unit | Control | Copper | P-Value<br>[Effect size] |
| --- | --- | --- | --- | --- |
| *Inter-bout Interval* |  |  |  |  |
| Vertical displacement | mm | 0.02 [0.62] | -0.32 [0.65] | **0.0473** [0.73] |
| Vertical velocity | mm s<sup>-1</sup> | -0.01 [0.30] | -0.27 [0.45] | **0.0183** [1.00] |
| Duration | s | 1.78 [2.29] | 1.42 [1.15] | **0.0300** [0.29] |
| Pitch | deg | 8.75 [17.74] | 12.88 [16.80] | **0.0473** [0.32] |
| Angular velocity | ° s<sup>-1</sup> | -2.70 [3.71] | -6.59 [7.00] | **0.0060** [1.06] |
| *Bout* |  |  |  |  |
| Peak speed | mm s<sup>-1</sup> | 11.48 [6.74] | 15.37 [10.36] | 0.0718 [0.67] |
| Trajectory | deg | 9.77 [29.33] | 13.53 [27.02] | 0.1481 [0.17] |
| Vertical displacement (climb) | mm | 0.37 [0.57] | 0.53 [0.83] | 0.0718 [0.35] |
| Vertical displacement (dive) | mm | -0.16 [0.23] | -0.21 [0.34] | **0.0473** [0.25] |
| Attack angle (climb) | deg | 3.76 [6.92] | 1.14 [6.59] | **0.0300** [0.52] |
| Attack angle (dive) | deg | -5.25 [5.37] | -6.27 [6.15] | 0.1050 [0.25] |
| Steering rotation gain^ |  | 0.69 [0.02] | 0.74 [0.02] | **0.0367** [1.58] |
| Righting rotation gain^ |  | 0.18 [0.01] | 0.18 [0.02] | 0.6281 [0.31] |
| Fraction of climbs | % | 66 [4] | 75 [7] | 0.0718 [1.59] |
| Fraction of dives | % | 32 [4] | 23 [7] | 0.0718 [1.76] |

**Table S3:** Inter-bout interval vertical velocity by time post-treatment. Early and late represent data collected in the first and second 5 hours after the treatment, respectively, for each 24-hour behavior session using Dark-Dark dataset. P-value calculated using Wilcoxon rank sum (*α* = 0.05).

| Treatment | Time<br>Post-Treatment | Median<br>[IQR] | P-Value |
| --- | --- | --- | --- |
| Control | Early | -0.06 [0.29] | 0.14164 |
|  | Late | 0.02 [0.45] |  |
| Copper | Early | -0.3 [0.20] | 0.59081 |
|  | Late | -0.29 [0.41] |  |

**Table S4:** Number of inter-bout intervals and bouts across datasets. Percentages reflect median fraction and [IQR] of total bouts for that group and time point. P-value calculated using Mann-Whitney U test with *α* = 0.05 for dark and light datasets, and using one-way ANOVA for longitudinal data. Bold values indicate significance.

| Dataset | Condition | IBIs | Climb bouts | Flat bouts | Dive bouts | Total bouts | Climb %<br>P-Value |
| --- | --- | --- | --- | --- | --- | --- | --- |
| Dark-Dark | control | 31228 | 22537 (66 [4]%) | 681 (2 [0.4]%) | 10823 (32 [4]%) | 34041 | 0.0718 |
|  | copper | 14407 | 10811 (75 [7]%) | 258 (2 [0.5]%) | 3630 (23 [7]%) | 14699 |  |
| Light-Dark | control | 174820 | 81680 (46 [5]%) | 5394 (3 [0.4]%) | 95460 (51 [5]%) | 182534 | **0.0013** |
|  | copper | 120012 | 75311 (60 [13]%) | 3165 (3 [0.4]%) | 48602 (38 [13]%) | 127078 |  |
| *Longitudinal—* | control | 21526 | 16150 (69 [12]%) | 544 (2 [0.9]%) | 7138 (29 [11]%) | 23832 | 0.4717 |
| Early Phase | copperEarly | 18504 | 12205 (68 [8]%) | 371 (2 [1]%) | 5826 (30 [7]%) | 18402 |  |
|  | copperLate | 30536 | 21072 (62 [9]%) | 811 (3 [1]%) | 11159 (34%) | 33042 |  |
| *Longitudinal—* | control | 17020 | 12149 (71 [7]%) | 202 (1 [0.2]%) | 4831 (27 [7]%) | 17182 | 0.6314 |
| Late Phase | copperEarly | 15902 | 11270 (68 [2]%) | 264 (1 [1]%) | 5093 (31 [2]%) | 16627 |  |
|  | copperLate | 19892 | 12242 (71 [20]%) | 231 (1 [0.9]%) | 5683 (27 [19]%) | 18156 |  |

**Table S5:** Effects of vision on swimming after lateral line loss. Median [IQR] used for non-normal distributions of raw data. N = 150 LD control, 150 LD copper. P-value calculated using two-way ANOVA and Tukey’s HSD post-hoc comparison (*α* = 0.05; bold numbers indicate significance), and effect size calculated using Cohen’s d. IBI, inter-bout interval

| Parameter | Unit | LD Control | LD Copper | Effect of Light | Effect of Treatment | Interaction Effect |
| --- | --- | --- | --- | --- | --- | --- |
| IBI Duration | s | 0.69 [0.54] | 0.63 [0.50] | **8.4e-13** | **9.6e-3** | **9.5e-3** |
| Vertical acceleration | mm s<sup>-2</sup> | -0.04 [0.03] | -0.16 [0.08] | **0.0216** | **1.5e-5** | 0.9176 |
| Bout trajectory | deg | -1.29 [18.31] | 4.28 [21.82] | **7.4e-8** | **1.17e-3** | 0.4744 |

**Table S6:**
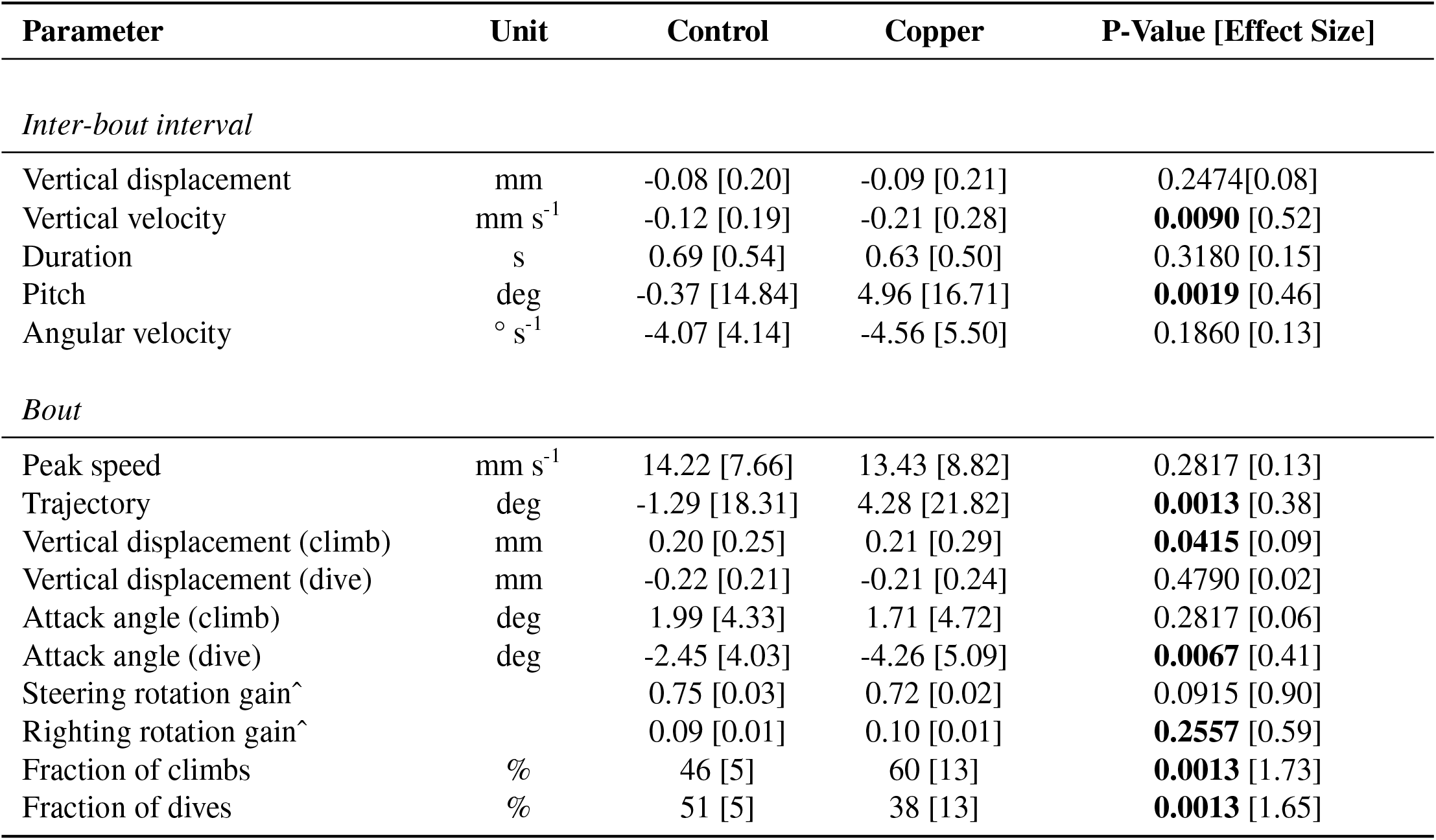
Effects of lateral line loss on swimming in the light. Median [IQR] used for non-normal distributions of raw data. N = 150 LD control, 150 LD copper. P-value calculated using Mann-Whitney U test (*α* = 0.05; bold numbers indicate significance), and effect size calculated using Cohen’s d. ^Steering and righting rotation gain p-value calculated using unpaired t-test (mean and standard deviation listed).

| Parameter | Unit | Control | Copper | P-Value [Effect Size] |
| --- | --- | --- | --- | --- |
| *Inter-bout interval* |  |  |  |  |
| Vertical displacement | mm | -0.08 [0.20] | -0.09 [0.21] | 0.2474[0.08] |
| Vertical velocity | mm s<sup>-1</sup> | -0.12 [0.19] | -0.21 [0.28] | **0.0090** [0.52] |
| Duration | s | 0.69 [0.54] | 0.63 [0.50] | 0.3180 [0.15] |
| Pitch | deg | -0.37 [14.84] | 4.96 [16.71] | **0.0019** [0.46] |
| Angular velocity | ° s<sup>-1</sup> | -4.07 [4.14] | -4.56 [5.50] | 0.1860 [0.13] |
| *Bout* |  |  |  |  |
| Peak speed | mm s<sup>-1</sup> | 14.22 [7.66] | 13.43 [8.82] | 0.2817 [0.13] |
| Trajectory | deg | -1.29 [18.31] | 4.28 [21.82] | **0.0013** [0.38] |
| Vertical displacement (climb) | mm | 0.20 [0.25] | 0.21 [0.29] | **0.0415** [0.09] |
| Vertical displacement (dive) | mm | -0.22 [0.21] | -0.21 [0.24] | 0.4790 [0.02] |
| Attack angle (climb) | deg | 1.99 [4.33] | 1.71 [4.72] | 0.2817 [0.06] |
| Attack angle (dive) | deg | -2.45 [4.03] | -4.26 [5.09] | **0.0067** [0.41] |
| Steering rotation gain^ |  | 0.75 [0.03] | 0.72 [0.02] | 0.0915 [0.90] |
| Righting rotation gain^ |  | 0.09 [0.01] | 0.10 [0.01] | **0.2557** [0.59] |
| Fraction of climbs | % | 46 [5] | 60 [13] | **0.0013** [1.73] |
| Fraction of dives | % | 51 [5] | 38 [13] | **0.0013** [1.65] |

**Table S7:** Effects of hair cell regeneration after lateral line ablation. Median [IQR] used for non-normal distributions of raw data. N=85 control larvae, 94 larvae treated with copper in the early phase (7–8 dpf), and 64 larvae treated with copper in the late phase (14–15 dpf). Data from late phase of recordings. P-value calculated using one-way ANOVA with Tukey’s HSD post-hoc comparison (*α* = 0.05; bold numbers indicate significance) and effect size calculated using Cohen’s d and enclosed in []. ^Steering and righting gain p-value calculated using unpaired t-test (mean and standard deviation listed).

| Parameter | Unit | Control | Copper Early | Copper Late | One-Way ANOVA P-Value | Control v. Early Post-Hoc | Early v. Late Post-Hoc | Control v. Late Post-Hoc |
| --- | --- | --- | --- | --- | --- | --- | --- | --- |
| *Inter-bout Interval* |  |  |  |  |  |  |  |  |
| Vertical displacement | mm | -0.05 [0.79] | -0.04 [0.70] | -0.35 [0.89] | **0.0048** | 0.9000[0.01] | **0.0122** [0.51] | **0.0095** [0.47] |
| Vertical velocity | mm s<sup>-1</sup> | -0.04 [0.31] | -0.03 [0.24] | -0.23 [0.46] | **0.0004** | 0.9000 [0.03] | **0.0012** [0.68] | **0.0013** [0.63] |
| Duration | s | 2.50 [3.69] | 2.92 [3.49] | 1.86 [2.48] | 0.1832 | 0.7858 [0.18] | 0.1671 [0.53] | 0.4424 [0.32] |
| Pitch | deg | 11.25 [21.51] | 7.08 [18.09] | 9.57 [19.11] | 0.5480 | 0.5569 [0.28] | 0.9000 [0.18] | 0.6640 [0.11] |
| Angular velocity | ° s<sup>-1</sup> | -2.63 [4.77] | -3.02 [4.57] | -5.60 [7.46] | **0.0237** | 0.9000 [0.11] | **0.0476** [0.55] | **0.0386** [0.63] |
| *Bout* |  |  |  |  |  |  |  |  |
| Peak speed | mm s<sup>-1</sup> | 10.46 [6.56] | 9.88 [5.89] | 13.07 [9.91] | **0.0180** | 0.3964 [0.13] | **0.0119** [0.55] | **0.0330** [0.43] |
| Trajectory | deg | 17.82 [38.01] | 12.52 [35.13] | 14.50 [37.83] | 0.6417 | 0.4374 [0.19] | 0.2474 [0.07] | 0.2474 [0.11] |
| Vertical displacement (climb) | mm | 1.20 [0.98] | 1.09 [0.90] | 1.52 [1.07] | 0.5238 | 0.3182 [0.15] | 0.0067 [0.57] | 0.0090 [0.41] |
| Vertical displacement (dive) | mm | -0.19 [0.39] | -0.14 [0.34] | -0.25 [0.46] | 0.2605 | 0.3964 [0.19] | 0.0946 [0.40] | 0.0639 [0.21] |
| Attack angle (climb) | deg | 9.77 [6.80] | 10.68 [6.21] | 7.55 [6.11] | 0.0671 | 0.1860 [0.18] | 0.0119 [0.70] | 0.0639 [0.46] |
| Attack angle (dive) | deg | -5.37 [6.00] | -4.33 [6.46] | -5.78 [6.19] | 0.3155 | 0.1860 [0.22] | 0.0517 [0.31] | 0.4790 [0.09] |
| Steering rotation gain^ |  | 0.67 [0.03] | 0.67 [0.06] | 0.67 [0.07] | 0.9937 | 0.8980 [0.06] | 0.9592 [0.02] | 0.9584 [0.02] |
| Righting rotation gain^ |  | 0.16 [0.03] | 0.16 [0.02] | 0.18 [0.01] | 0.4476 | 0.9026 [0.06] | 0.1924 [0.68] | 0.3251 [0.50] |
| Fraction of Climbs | % | 71 [7] | 68 [2] | 71 [20] | 0.6314 | 0.0946 [0.08] | 0.2474 [0.08] | 0.4790 [0.007] |
| Fraction of Dives | % | 27 [7] | 31 [2] | 27 [19] | 0.6440 | 0.0781 [0.08] | 0.2474 [0.08] | 0.4790 [0.007] |

